# Updated definitions on piezophily as suggested by hydrostatic pressure dependence on temperature

**DOI:** 10.1101/2020.08.31.275172

**Authors:** Alberto Scoma

**Affiliations:** Engineered Microbial Systems Laboratory (EMS-Lab), Biological and Chemical Engineering Section, Dept. of Engineering, Aarhus University, Hangøvej 2, 8200, Aarhus N, Denmark

## Abstract

Microbial preference for elevated hydrostatic pressure (HP) is a recognized key feature of environmental and industrial processes. HP effects on macromolecules and, consequently, cell functionality has been accurately described in the last decades. While there is little debate about the importance of HP in shaping microbial life, a systematic definition of microbial preference for increased HP is missing. The lack of a consensus about ‘true’ piezophiles, and ‘low’ or ‘high’ HP levels, has deleterious repercussions on microbiology and biotechnology. As certain levels are considered ‘low’ they are not applied to assess microbial activity. Most microorganisms collected in deep waters or sediments have not been tested (nor isolated) using the corresponding HP at which they were captured. Microbial response to HP is notoriously dependent on other environmental parameters, most notably temperature, but also on availability of nutrients, growth substrate, pH and salinity. This implies that countless isolates retrieved from ambient pressure conditions may very well require increased HP to grow optimally, as already demonstrated in both Archaea and Bacteria.

In the present study, I collected the data from described piezophilic isolates and used the fundamental correlation existing between HP and temperature, as first suggested in seminal works by Yayanos, to update the definition of piezophiles. Thanks to the numerous new piezophilic isolates available since such seminal studies, the present analysis brings forward updated definitions which concern 1) the actual beginning of the piezosphere, the area in the deep sea where piezophiles thrive; 2) the HP thresholds which should be considered low, medium and high HP, and their implications for experimental design in Microbiology; and 3) the nature of obligate piezophiles and their location in the deep sea.

## Introduction

Hydrostatic pressure (HP) is the force exerted on an area by a fluid at rest. Its influence on macromolecules of biological relevance contributed to the evolution of life. In the biosphere, HPs from 0.1 to 200 MPa can affect intermolecular distances and conformation of polynucleotides (DNA and RNA), lipid bilayers and multimeric proteins (Oger and Jebbar 2010). Increased HP features today planktonic crustaceans, and marine invertebrates, fish, and deep-diving mammals, with Bacteria and Archaea able to thrive at tens of MPa (Bartlett 2002). Elevated HPs characterize the largest microbial ecosystems on Earth (*i.e*. the deep sea and subseafloor (Whitman et al 1998)), may feature extraterrestrial life (Bartlett 2002), and are key to the inactivation of pathogens and viruses in food and medicine production (Aertsen et al 2009). Yet, a systematic definition of microbial preference for increased HP is missing. Such preference is generally referred to as piezophily, descriptive of microorganisms growing better at increased HP, with obligate piezophiles (or hyperpiezophiles) unable to grow at ambient pressure, and piezosensitive strains growing best at ambient pressure. The lack of consensus about ‘true’ piezophiles extends to the formalization of ‘low’ or ‘high’ HP levels and to the actual beginning of the piezosphere, the area of the deep sea where piezophiles thrive.

HP linearly increases by 1 MPa every 100 m of seawater depth, and about three times as much in sediments (Oger and Jebbar 2010). Enhanced growth at elevated HP has been used to locate the beginning of the piezosphere at corresponding depths (*e.g*. 1’000 m below seawater level [bswl] if HP was equal to the corresponding ~10 MPa). However, different HP thresholds were proposed for both piezophiles and hyperpiezophiles: ≥10 MPa (Jannasch and Taylor 1984)); ≥10 MPa for piezophiles and ≥50 MPa for hyperpiezophiles (Fang et al 2010); >0.1 MPa for piezophiles and ≥60 MPa for hyperpiezophiles (Yayanos 1995)), but indicating moderate piezophily if HP optimum [HP_opt_] of an isolate under investigation was between 10 and 30 MPa (Eloe et al 2008); ≥40 MPa (as the average depth of the oceans is about 3750 m, (Horikoshi and Bull 2011)). Following deep-sea sampling, isolation and lab-scale testing, a seminal work by Yayanos in 1986 proposed that true piezophiles populated water depths of at least 2’000 m (≥20 MPa). A most significant concept introduced by Yayanos was that of PTk diagrams, where growth rates are plotted in three dimensional graphs versus HP and temperature (T). Such diagrams indicate that maximum growth rate [μ_max_] at HP_opt_ is observed when concomitantly adjusting T to an optimal value [T_opt_]. However, Yayanos work was almost exclusively based on psychrophiles (microorganisms with a T_opt_ ≤15°C). The isolation of several new piezophiles in recent years, particularly with T_opt_ >15 °C, is an opportunity to revise the role of T on piezophiles. Deep-sea environments exposed to elevated HPs commonly experience low T (*i.e*. 2 °C). Exceptions are the deep, warm Sulu (~10 °C), Mediterranean (~13.5 °C) and Red sea (~20 °C); deep hydrothermal vents, where microorganisms may grow at >110 °C; and deep sub-seafloors where T increases 25 °C every km underground (Jebbar et al 2015, Yayanos 1995). Notwithstanding the many HP-T combinations in the environment, this correlation has not been systematically addressed.

The most commonly accepted definition for piezophily is physiological, and states that piezophiles show a μ_max_ above the atmospheric pressure of 0.1 MPa. This straightforward definition, however, does not appropriately describe microbial preference for elevated HPs. Can a microorganism with a HP_opt_ of 5 MPa be regarded as piezophilic as one with a HP_opt_ of 50 MPa? If intermediate HP values distinguish ‘moderate’ and ‘true’ piezophiles, they should be appropriately assessed *e.g*. by cultivation.

Cultivation is however a complicated task. Maximum cell density and μ_max_ may occur at different HPs. (Martini et al 2013) proposed a cross coefficient to describe piezophily including both such factors. Although this cross coefficient has not been adopted in later reports, the study has merit in highlighting the need for proper classification of piezophiles. Furthermore, aside T (Yayanos 1986), microbial preference for increased HP may also depend on availability of nutrients (Jannasch and Wirsen 1984), growth substrate (Jannasch and Taylor 1984, Yayanos and Chastain 1999), pH (Matsumura et al 1974) and salinity (Harrison et al 2013). All such dependencies are seldom tested on new piezophilic isolates.

The absence of an exhaustive and commonly accepted definition of microbial preference for HP has deep implications for the microbiology and biotechnology of piezophiles. As certain HP levels may be generally considered low, they are not applied to assess microbial activity. Most microorganisms collected underwater have not been tested (nor isolated) using the corresponding HP at which they were captured. The fact that several cultivation parameters can influence the microbial response to HP implies that several microorganisms originally collected from surface waters may have their HP_opt_ above atmospheric pressure. This was already demonstrated in both Archaea (*Methanococcus thermolithotrophicus* (Bernhardt et al 1988)) and Bacteria (*Clostridium paradoxum* (Scoma et al 2019)), where true μ_max_ at increased HP were determined by concomitantly varying T. At the time of writing the present note, there are only 86 documented microbial isolates with a HP_opt_ >0.1 MPa, an incredibly small group for a condition featuring the largest reservoir of prokaryotes on the planet (Whitman et al 1998). This lack of information has the worst implications when assessing the activity of microorganisms collected at shallower depths in the oceans (*e.g*. from meso- and bathypelagic waters), where hydrological conditions favor the presence of both HP-adapted (autochthonous) and HP-sensitive communities (allochthonous, *e.g*. sinking from surface waters), as opposed to deep, stratified waters (Tamburini et al 2013).

In the present note, I have collected data describing known piezophiles whose preference for increased HP has been assessed in laboratory tests, and discussed such preference based on their T_opt_. The objective was to improve the standing definition of piezophily relying on the existing dataset of cultured isolates, and individuate whether intermediate HPs exist to distinguish ‘moderate’ from ‘true’ piezophiles, indicative of the beginning of the piezosphere. Even though cultivation-based definitions may be biased by microbial cultivability, with only a minority of the existing microorganisms in the cultivable fraction, characterization of available isolates is the only way to learn the true HP ranges for a particular species.

## Results

The 86 described piezophiles (HP_opt_ >0.1) were grouped according to their T preference in piezopsychro- (T_opt_ ≤15 °C), piezomeso- (16< T_opt_ <49 °C) and piezothermophiles (T_opt_ ≥50 °C). There are currently 48, 17 and 21 prokaryotic species of documented piezopsychro-, piezomeso- and piezothermophilic nature, 71 of which are Bacteria and 15 Archaea (Table 1).

### Growth rates

All μ_max_ values in described piezophiles were plotted versus either HP_opt_ (Fig. S1A) or T_opt_ (Fig. S1B) to explain growth based on HP or T independently of one another. High HP_opt_ is consistent with low μ_max_ while the opposite is true for T_opt_. Aside these general observations, no strict correlation appears evident. Another approach was to divide piezophiles for T preference but neglect HP_opt_. The average μ_max_ in piezopsychro-, piezomeso- and piezothermophiles increases with T, namely 0.22±0.13, 0.42±0.38 and 1.25±0.84 h^−1^ (Table 1). However, only the average μ_max_ in piezothermophiles is significantly different from the other groups (t-test, *p*< 0.0011), and there is no difference between piezopsychro- and piezomesophiles (*p*>0.05). The opposite approach (*i.e*. neglecting T classification) was attempted using HP thresholds as suggested by (Yayanos 1995): piezophiles >0.1 MPa up to 60 MPa; and hyperpiezophiles >60 MPa. Average μ_max_ was comparable (*p*>0.05, 0.55±0.61 and 0.41±0.69 h^−1^, respectively). Overall, neither HP_opt_ nor T_opt_ alone appropriately describe the differences in piezophiles.

The combined effect of HP and T on growth was then assessed. The HP_opt_ and T_opt_ of every piezophile was ‘normalized’ diving each by their corresponding μ_max_ (Fig. 1), thus expressing the rate at which either HP_opt_ or T_opt_ increases with respect to their μ_max_. These rates were exponentially correlated in piezopsychro-, piezomeso- and piezothermophiles (r^2^ equal to 0.62 (n=37), 0.78 (n=12) and 0.58 (n=17), respectively). The regression line indicates that to attain μ_max_ HP and T must be combined in a specific, optimal proportion. Suboptimal combinations of HP and T not resulting into μ_max_ plot far from the regression line (*i.e*. have a lower μ). The relevance of HP-T correlations is only observed when separating piezophiles in the three T groups, as it can be inferred by the large variations in the axes of Fig. 1. Two data points were omitted from piezopsychrophiles, namely *Profundimonas piezophila* YC-1 and *Rhodobacterales bacterium* PRT1. These are the two slowest piezophiles isolated so far (μ_max_ <0.02 h^−1^), therefore normalized HP_opt_ and T_opt_ resulted out of scale. However, their inclusion would increase r^2^ from 0.62 to 0.87. One data point was removed from piezothermophiles for the same reason (*Archaeoglobus fulgidus* VC-16^T^), its inclusion slightly reducing r^2^ from 0.58 to 0.54.

**Figure 1.**
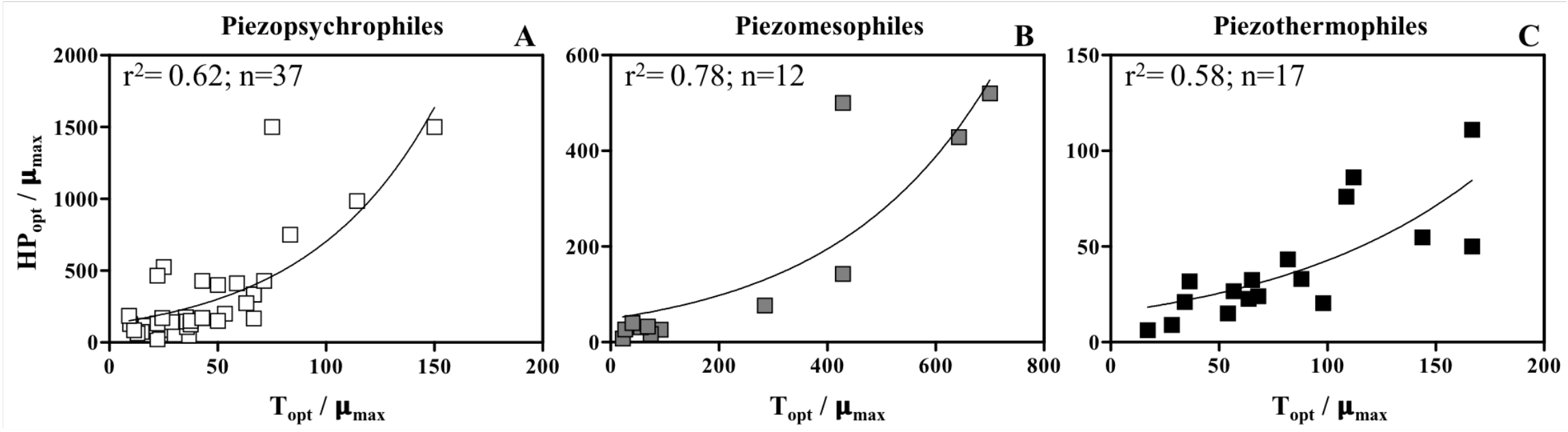
Correlation between the rate increase in HP_opt_ (HP_opt_/μ_max_) and rate increase in T_opt_ (T_opt_/μ_max_) with respect to μ_max_ in the piezopsychro-, piezomeso- and piezothermophiles. Statistical correlation was obtained with GraphPad Prism 5, nonlinear regression, exponential growth equation, least square (ordinary) fit.

### Capture depth and HP_opt_

The relationship between piezophiles’ capture depth and HP_opt_ is reported in Fig. 2A, while that between HP_opt_ and T_opt_ in Fig. 2B. There is an almost linear correlation between capture depth and HP_opt_ in piezopsychrophiles (r^2^=0.69, n=48). This correlation is irrelevant in piezomeso- (r^2^=0.04, n=15) and piezothermophiles (r^2^=0.06, n=21). Isolates with a markedly lower HP_opt_ than that experienced at capture depth may indicate they did not originally belong in that environment (*i.e*. they were allochthonous). As HP increase differs underground, two piezomesophiles collected at about 2 km in the deep subseafloor (Fang et al 2017) were removed when assessing linear correlation, all others were included. It may be concluded that capture depth in the water column is seldom (if at all) relevant except for piezopsychrophiles. Removal of the six Bacteria from piezothermophiles to identify a correlation only in Archaea had just a slight effect on the goodness of fit (from 0.06 to 0.09).

**Figure 2.**
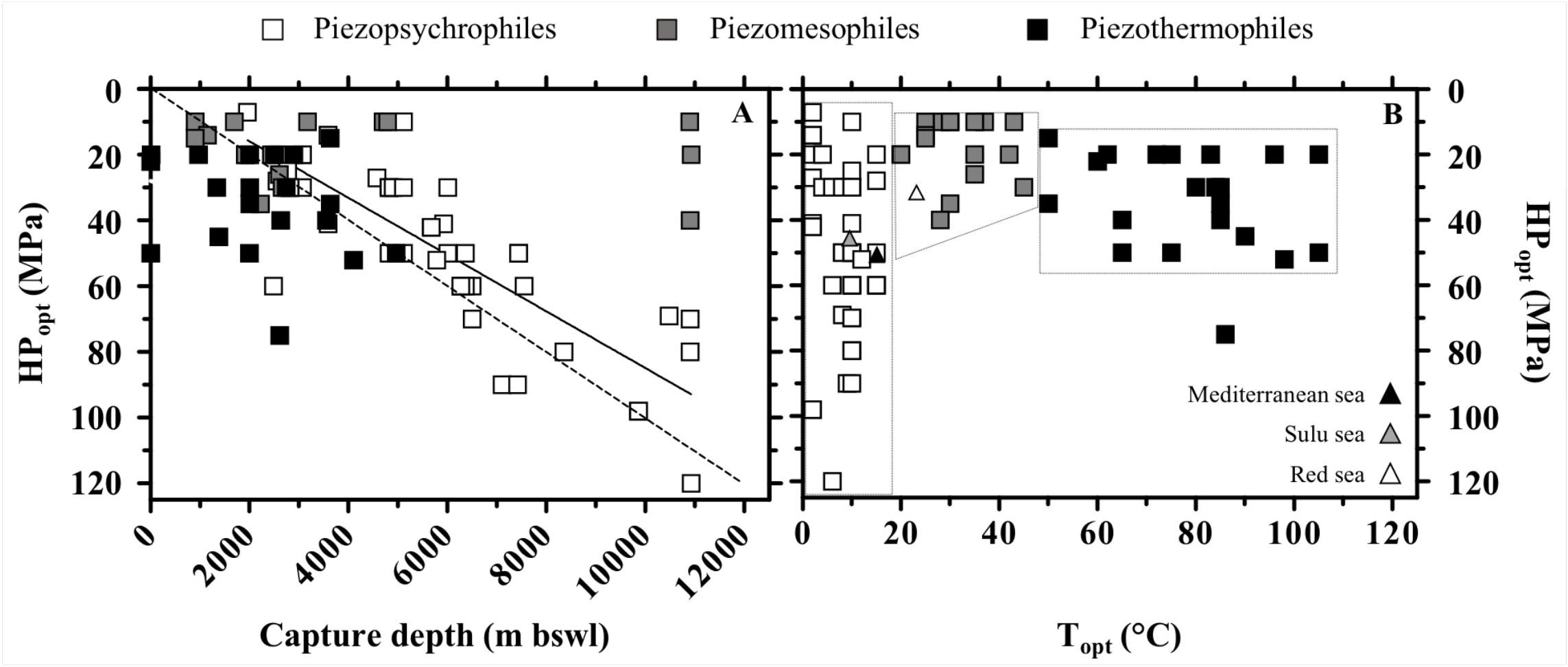
Correlation between optimal HP (HP_opt_) and capture depth (A) and between HP_opt_ and optimal temperature (T_opt_) (B) in described piezophilic isolates. In Fig (A): the straight line indicates the linear regression for piezopsychrophiles (r^2^=0.69, n=48); the dotted line indicates the linear increase of HP with increasing depth in seawater. In Fig (B): approximate maximum T at the deepest point (expressed as equivalent HP) in the warmest seas in the world; dotted quadrilaterals define areas of HP_opt_-T_opt_ for known piezophiles (all data in Table 1). Keys reported in the graph. In Fig. (A) statistical correlation was obtained with GraphPad Prism 5, linear regression.

The reason why HP_opt_ in piezothermophiles does not correlate with HP at capture depth is unclear. Microbial dispersion from the deep hot biosphere may partially account for this (Hoshino et al 2017), however, most piezothermophilic isolates derive from hydrothermal vents. Resilience of sinking allochthonous microorganisms from warmer surface waters may represent a relevant bias in piezomesophiles. Five isolates show HP_opt_ remarkably lower compared to HP at capture depth (*Salinimonas sediminis* N102^T^, *Paraoceanicella profunda* D4M1, *Shewanella profunda* LT13a and *Pseudodesulfovibrio indicus* J2^T^ and *Muricauda hadalis* MT-229^T^; Table 1B). Their removal increases the goodness of fit from r^2^=0.04 to 0.53. However, such a selective data treatment may just confirm the lack of correlation between capture depth and HP_opt_ in piezomesophiles. A similar data refining for piezopsychrophiles (five strains) has no effect on the goodness of fit. The small discrepancy between linear HP increase with depth (dotted line, Fig. 2A) and linear regression of HP_opt_ with capture depth in piezopsychrophiles (straight line, Fig. 2A) indicates a slightly lower HP_opt_ in piezopsychrophiles as compared to HP at capture depth. This consolidates previous observations with single piezopsychrophiles (Deming et al 1984, Yayanos et al 1982).

### Minimum HP_opt_

HP_opt_ is the HP at which μ_max_ is observed in a piezophile. In particular, the lowest HP_opt_ values indicate that no piezophile was found growing optimally with less HP. Examination of the lowest HP_opt_ may thus be used to distinguish ‘low’ from ‘high’ HP.

There is presently a slight difference in the lowest HP_opt_ according to T preference: in piezopsychrophiles, 8 isolates have HP_opt_ around 10 to 20 MPa (with the sole exception of *Shewanella* sp. SC2A at 7 MPa, although its growth rate is almost identical up until 14 MPa, 0.076 *vs*. 0.072 h^−1^, respectively (Yayanos et al 1982)); in piezomesophiles, the lowest HP_opt_ is always 10 MPa; for piezothermophiles, the minimum is at 20 MPa with the sole exception of *Thioprofundum lithotrophica* 106 at 15 MPa. When piezophiles are considered altogether, the minimum HP_opt_ ranges between 10 and 20 MPa. In other words: i) at least 10 MPa are generally required to observe piezophily, and ii) ≥20 MPa piezophilic traits can be found in any T group.

Using minimum HP_opt_ in described piezophiles as a threshold to define true piezophily has several advantages: it is independent of sampling procedures and environment (*i.e*. issues with sample preservation in challenging environments; sampling of allochthonous microorganisms); given its cultivation-based nature, it can be reproduced in any laboratory, meaning that the scientific debate is open to a broader research community; optimization of culture conditions allows to update these thresholds in time. Nonetheless, the current trends may just reflect the lack of laboratory testing at very low HPs (*e.g*. 5 MPa) although some of the isolates showing such minimum HP_opt_ were also grown at lower HPs (12.5, 25 and 50% in piezopsychro-, piezomeso- and piezothermophiles, respectively). Besides, countless microorganisms isolated from surface water have never been tested for HP tolerance. This is particularly relevant for psychrophilic *Colwellia* and *Shewanella* collected from surface waters, as these genera have several members among the most extreme piezopsychrophiles (Table 1A); or for *Shewanella* in piezomesophiles (Table 1B).

**Table 1A.**
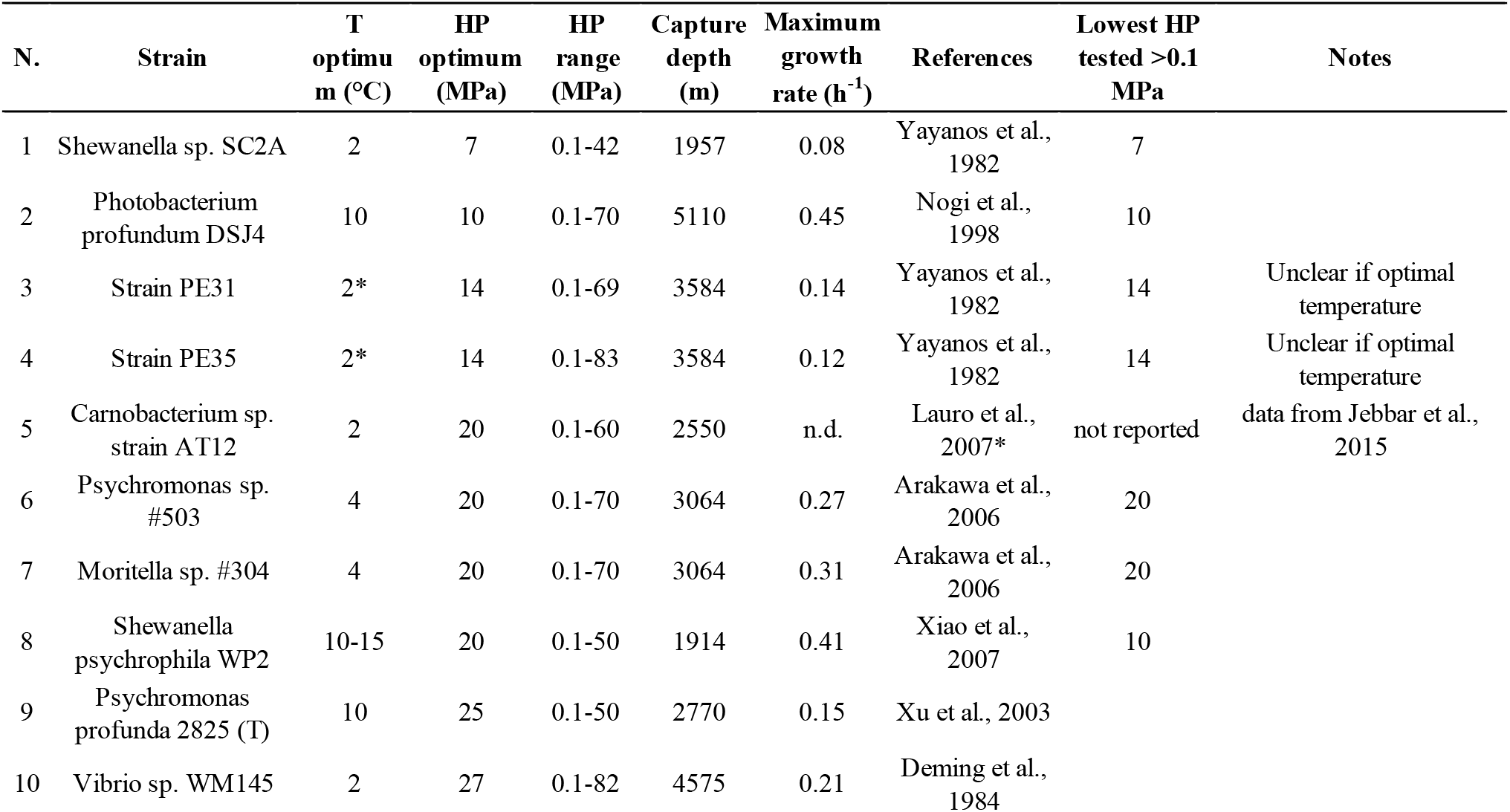
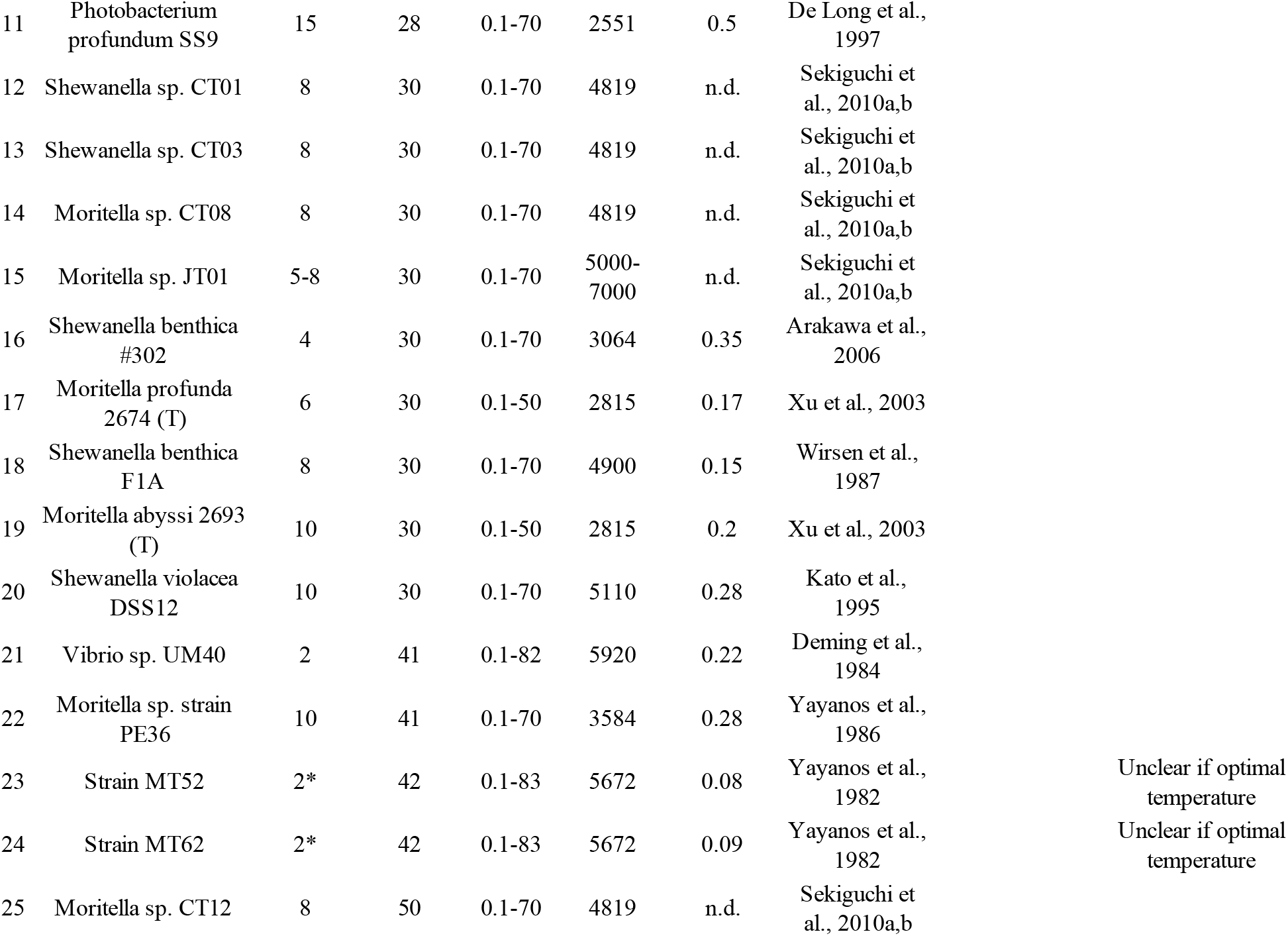
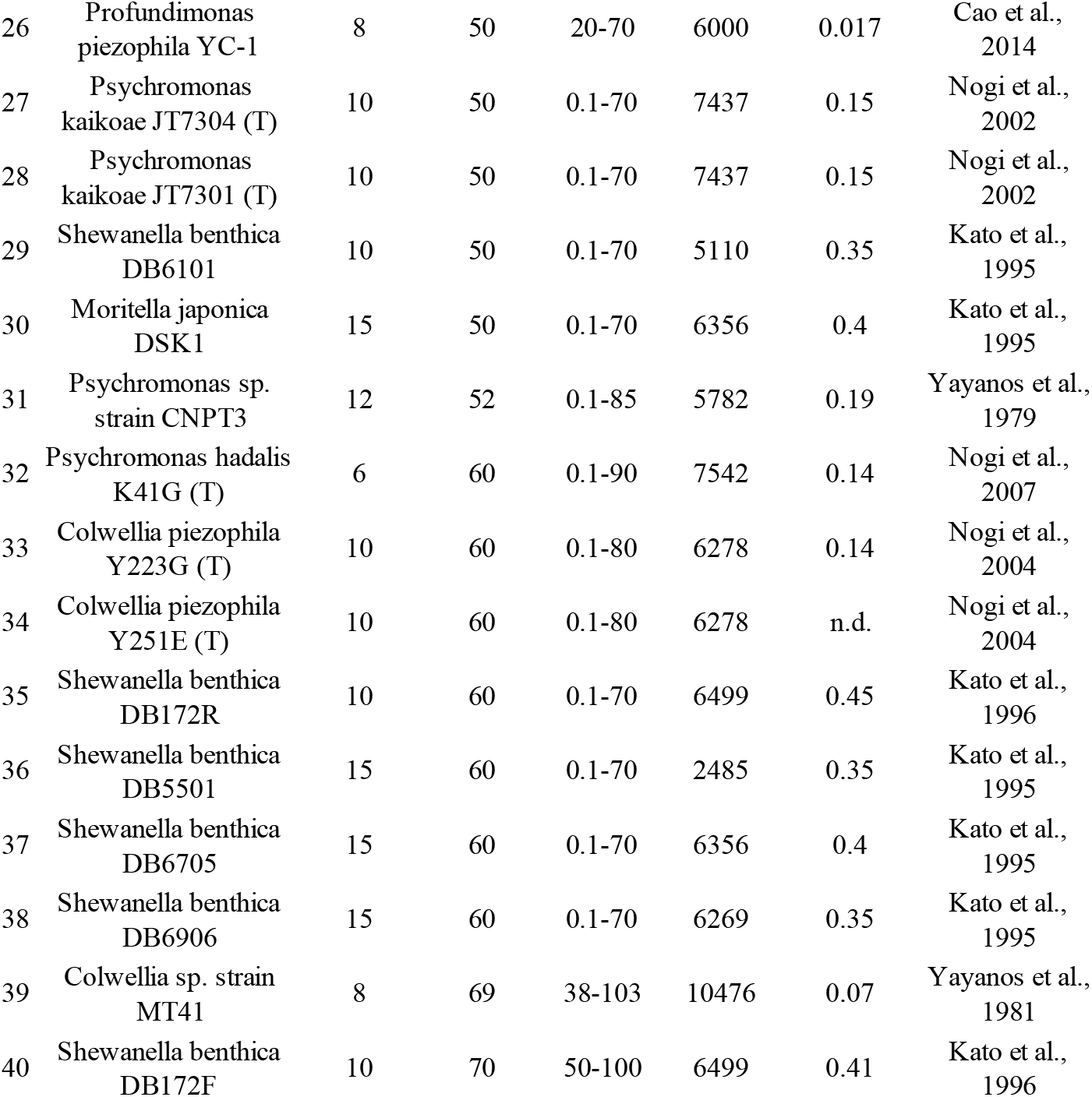
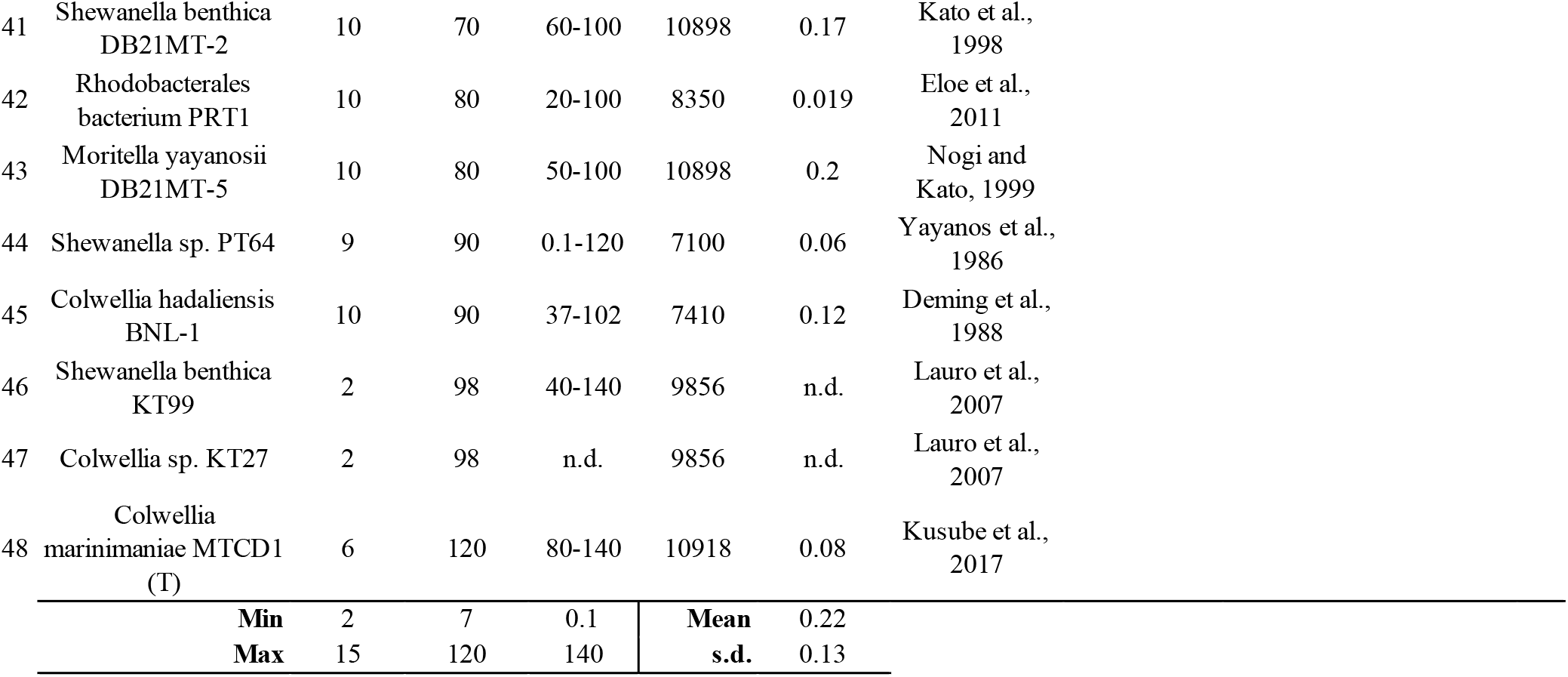
Described piezopsychrophilic isolates and their main features. Temperature boundaries were defined based on Morita RY (1975). Psychrophilic bacteria. Bacteriol Rev 39: 144-167. **Keys**: n.d., not determined; T, temperature; HP, hydrostatic pressure; (T), type strain. Asterisks (*) refer to their corresponding Note.

**Table 1B.**
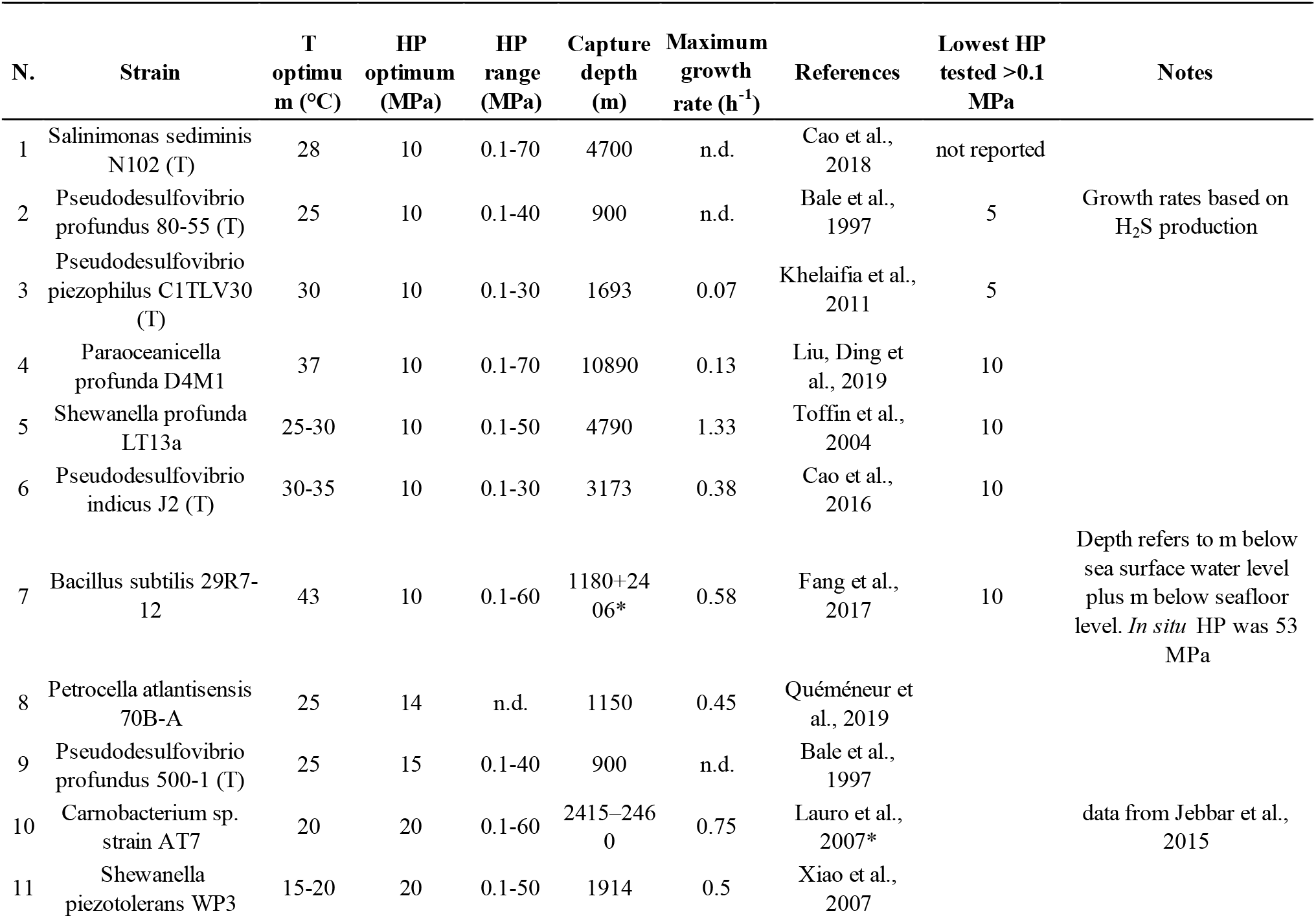
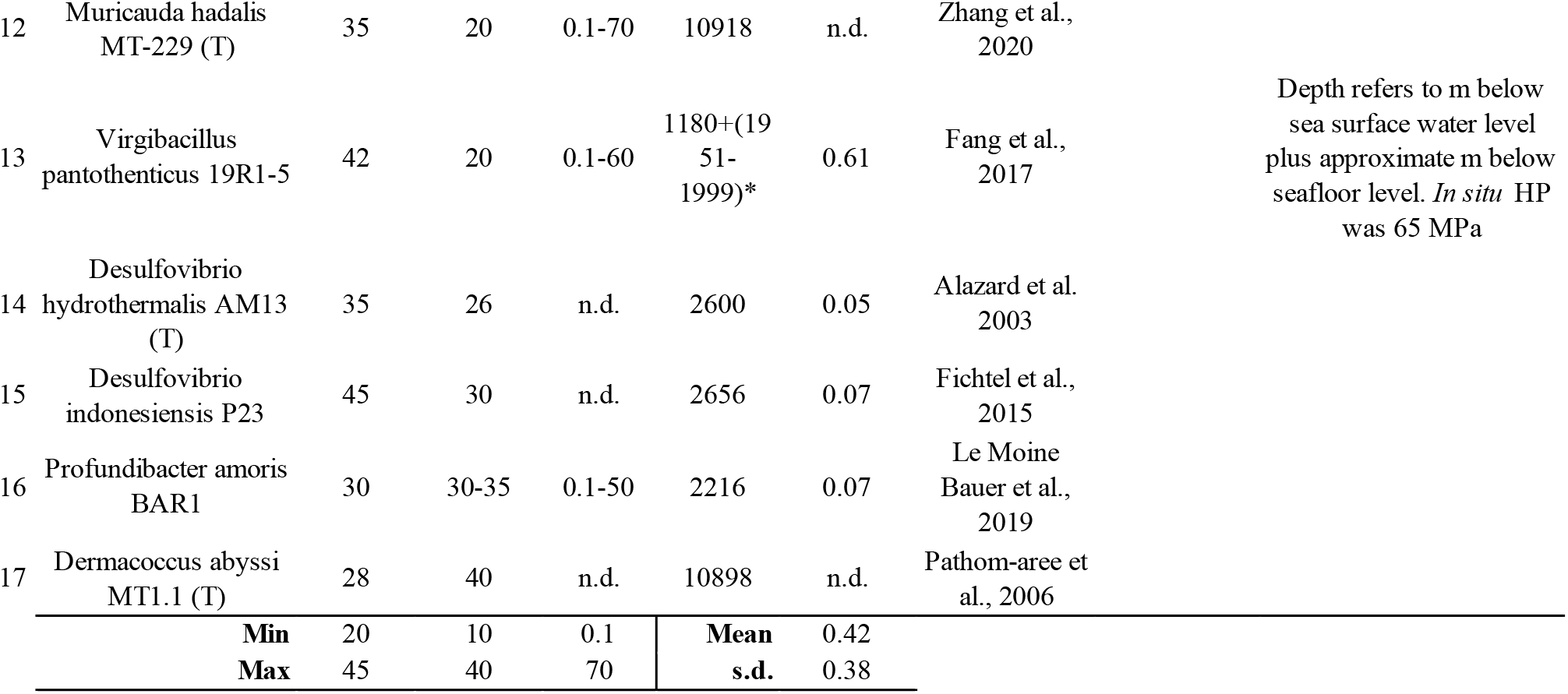
Described piezomesophilic isolates and their main features. Temperature boundaries were defined based on the definition of psychrophiles and thermophiles. **Keys**: n.d., not determined; T, temperature; HP, hydrostatic pressure; (T), type strain. Asterisks (*) refer to their corresponding Note.

**Table 1C.**
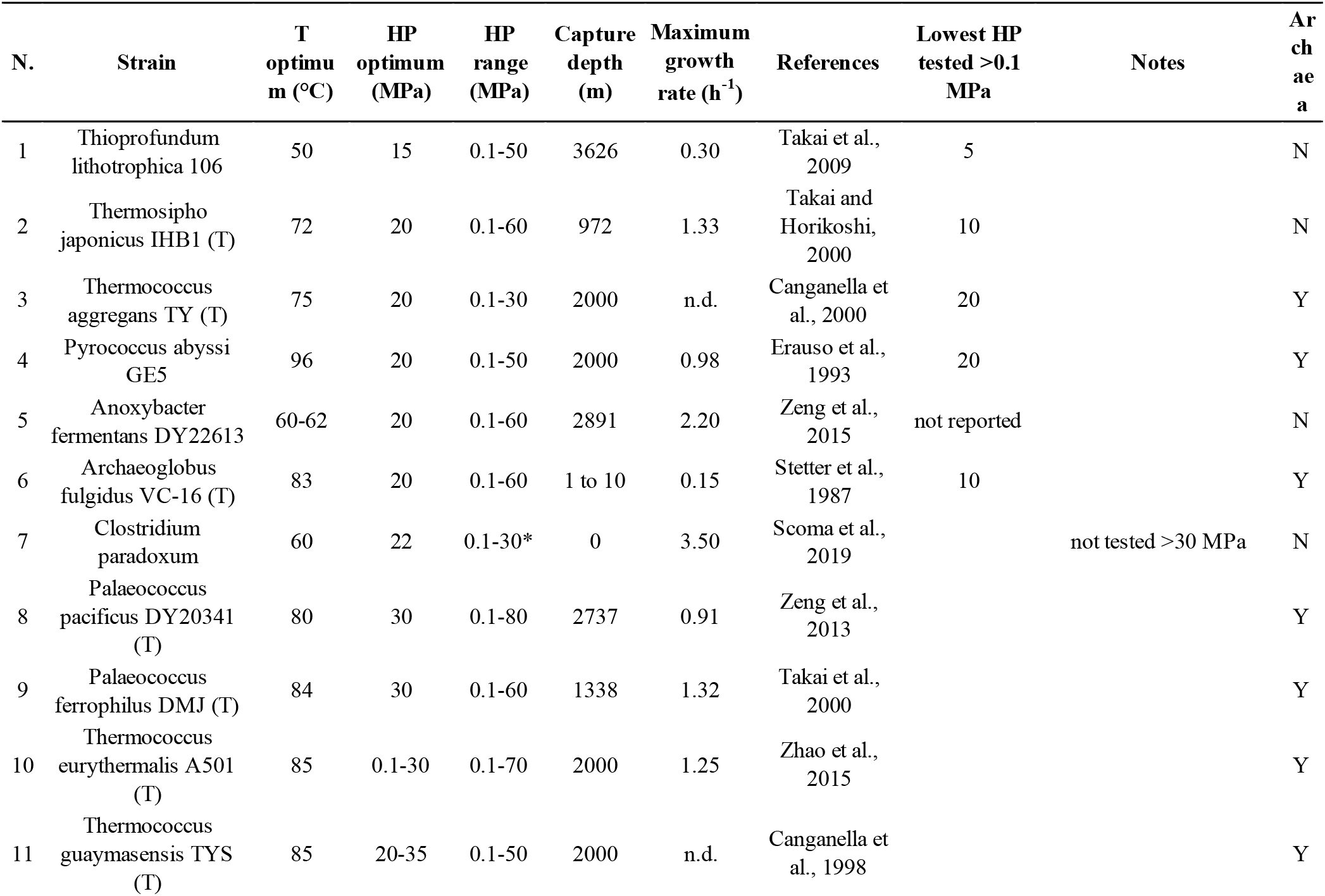
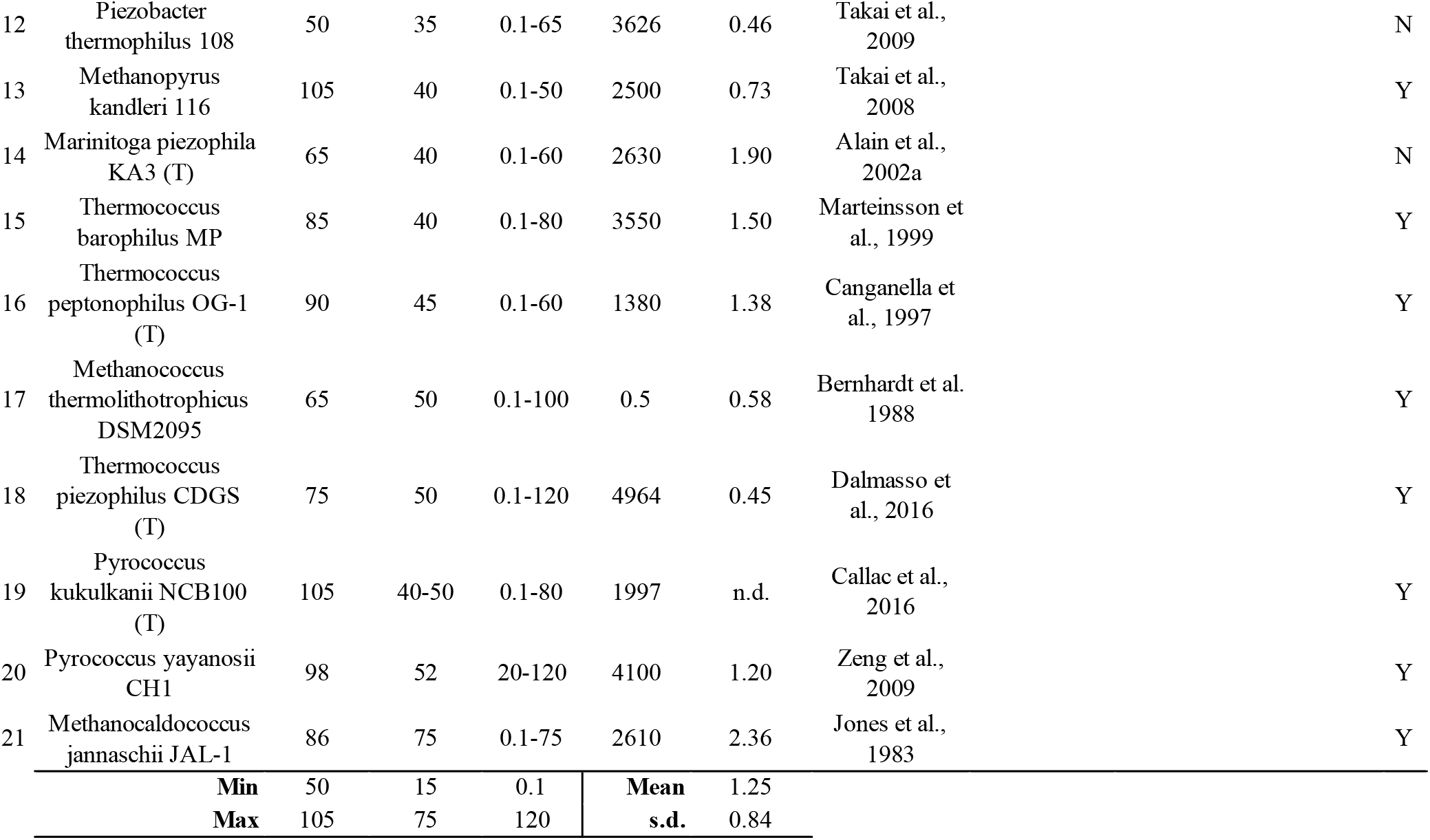
Described piezothermophilic isolates and their main features. Temperature boundaries were defined based on Zeikus JG (1979). Thermophilic Bacteria - Ecology, Physiology and Technology. Enzyme and Microbial Technology 1: 243-252. **Keys**: n.d., not determined; T, temperature; HP, hydrostatic pressure; (T), type strain; Y, yes; N, No. Asterisks (*) refer to their corresponding Note.

### T_opt_ and HP_opt_

Microbial preference for different T_opt_ and HP_opt_ defines three areas where described piezophiles grow optimally (Fig. 2B). Piezopsychrophiles, by definition growing in a narrow range of T, form a rectangle that stretches up to 120 MPa. Piezomesophiles grow in a quadrilateral area that currently lacks representatives at extreme T_opt_-HP_opt_ combinations. Piezothermophiles, typically growing within large T_opt_ variations, have an upper HP_opt_ not higher than 55 MPa, with one single exception (*Methanocaldococcus jannaschii* JAL-1).

Each of these areas is consistent with existing ecological niches. Cold, deep-oceans do experience the T-HP combinations which are optimal for the isolated piezopsychrophiles, a consideration which extends to piezothermophiles as hydrothermal vents are found at depths of ~5’000 m (Jebbar et al 2015). However, piezothermophiles are mostly Archaea (15/21, 71%), the most extreme bacterium represented by *Marinitoga piezophila* KA3^T^ (T_opt_ 65°C, HP_opt_ 40 MPa). Seafloor HP and T for the warmest deep seas on our planet (the Mediterranean, Sulu and Red sea) are indicated (Fig. 2B). These define an imaginary triangle which concerns piezopsychro- and piezomesophiles. The physiological properties of described piezomesophiles extend to T_opt_-HP_opt_ combinations well beyond this triangular area, particularly to higher T_opt_ which often involve anaerobic isolates collected from subseafloors (Table 1B). Nonetheless, no piezomesophile thus far has been isolated which requires the most extreme T_opt_-HP_opt_ combinations *e.g*. 45 °C and 50 MPa. Recently *Virgibacillus pantothenticus* 19R1-5 and *Bacillus subtilis* 29R7-12 were isolated from deep-subseafloor samples (between 1.9 and 2.5 km below seafloor level) characterized by such high T and HP (Fang et al 2017). These have T_opt_ close to T at capture depth (42-43 °C), but considerably lower HP_opt_ (20 and 10 MPa; while *in situ* levels were 53 and 65 MPa, respectively) (Fang et al 2017)). The difficulty in collection, preservation and cultivation of these samples represents a critical limit for isolation of extreme piezomesophiles.

### Hyperpiezophiles

While there are several obligate piezopsychrophiles (9/48), there is none for piezomesophiles, and just one for piezothermophiles (*Pyrococcus yayanosii* CH1) (Table 1). The term hyperpiezophiles thus currently describes a group of psychrophiles. Their lowest HP_opt_ is generally ~70 MPa (4/9 strains), with the exception of *P. piezophila* YC-1 at 50 MPa. The shallowest capture depth ranges between 6’000-6’500 m bswl.

## Discussion

The most widely shared definition of microbial preference for increased HP states that a microorganism is piezophilic when its μ_max_ is obtained at HPs >0.1 MPa. By setting the threshold to such a low reference value, this definition neglects the large variation in HP preference that exists among described piezophiles and the differential effects enhanced HP may impose on the vast diversity of microbial processes in nature. An additional cultivation-based approach is required to identify the threshold level above which HP-adapted microorganisms clearly separate from those thriving at ambient pressure.

In the present meta-analysis, the relevance of HP-T combinations first described by Yayanos (1986) was updated to include all currently described piezophiles, which were classified according to their T preference in piezopsychro-, piezomeso- and piezothermophiles. T preference appears to define piezophiles in several regards, namely: 1) μ_max_ is attained at specific HP-T combinations; the rates of both HP_opt_ and T_opt_ increase with respect to μ_max_ are exponentially correlated, and such correlation differs in the three T groups; 2) the concept of capture depth and its relation to HP_opt_ is only relevant in piezopsychrophiles; 3) piezopsychrophiles have a HP_opt_ slightly lower than the HP at capture depth; 4) there are no extreme piezomesophiles, perhaps an indication of the technical limitations in their collection and cultivation; 5) similarly, there are no obligate piezomesophiles; 6) obligate piezophily features piezopsychrophiles almost exclusively, with a minimum HP_opt_ of 70 MPa and capture depth of at least 6’000-6’500 m bswl; 7) HP_opt_ >75 MPa exclusively belong to piezopsychrophiles.

### Updated definitions

Based on the present meta-analysis, the following updated definitions are proposed:

1. As capture depth is only linearly correlated with HP_opt_ in piezopsychrophiles, and it complicates the comparison of isolates that present-day technology allows to collect from deep subseafloors, the definition of piezophily and piezosphere should refer to HP at capture depth (in MPa) rather than meters below sea or seafloor level.
2. Minimum HP_opt_ may be considered to define low, medium and high HP levels, *e.g*. when assessing microbial activity with environmental samples in laboratories or ship boards using samples carrying an unknown mixture of autochthonous and allochthonous microorganisms. Low HPs may be considered as those below which piezophiles across all T levels seldom have their HP_opt_; high HPs those above which piezophiles typically have their HP_opt_ irrespective of their T preference; and mild HP levels those in between these two extremes. The present data across all available prokaryotic isolates (n=86) suggests these thresholds to be: low <10 MPa; mild ≥10 and <20 MPa; and high ≥20 MPa.
3. As a consequence of point 2, moderate piezophiles may be defined as those with HP_opt_ <20 MPa, and true piezophiles those with HP_opt_ ≥20 MPa. The piezosphere would therefore begin at 20 MPa.
4. True hyperpiezophiles are so far almost exclusively psychrophilic in nature. Until isolation of a consistent number of piezomeso- and piezothermophiles indicates otherwise, true hyperpiezophiles are captured at least at 6’000-6’500 m bswl and have HP_opt_ ≥70 MPa.

## Acknowledgements

I am grateful to Prof. Kasper Urup Kjeldsen and Dr. Grégoire Michoud for the scientific discussions.

