## Supplementary material for "Updated definitions on piezophily as suggested by hydrostatic pressure dependence on temperature": Figure S1

**Figure S1. Correlation between maximum growth rates ($\mu_{max}$) and optimal hydrostatic pressure ($HP_{opt}$) or optimal temperature ($T_{opt}$) in piezophilic isolates**

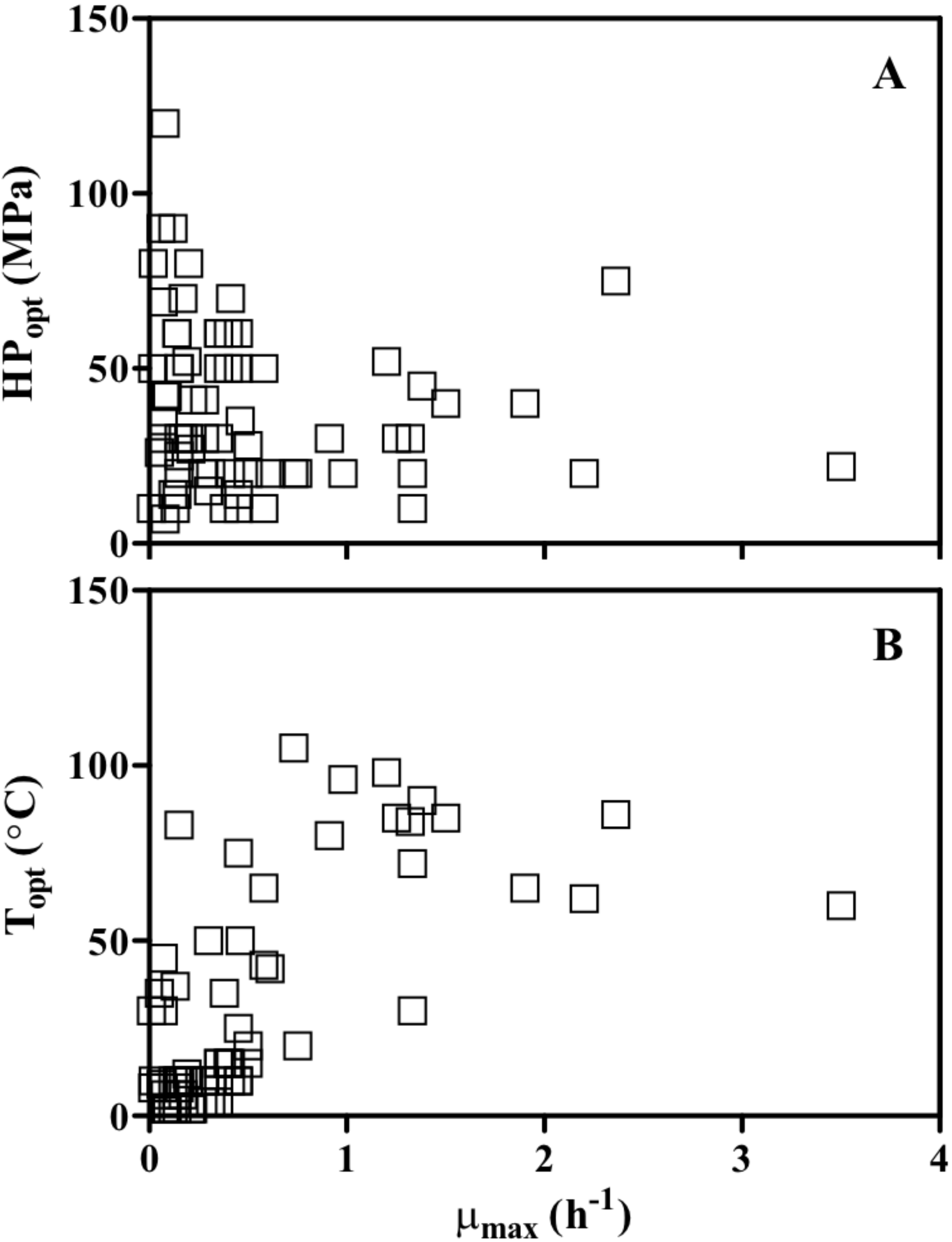
